# Whole blood transcriptome analysis reveals footprints of cattle adaptation to sub-arctic conditions

**DOI:** 10.1101/379925

**Authors:** Kisun Pokharel, Melak Weldenegodguad, Ruslan Popov, Mervi Honkatukia, Hanna Huuki, Heli Lindeberg, Jaana Peippo, Tiina Reilas, Stepan Zarovnyaev, Juha Kantanen

## Abstract

Indigenous cattle breeds in northern Eurasia have adapted to harsh climate conditions. The local breeds are important genetic resources with cultural and historical heritages, and therefore, their preservation and genetic characterization are important. In this study, we aim to identify genes and biological processes that are important for their adaptation to the cold and harsh conditions. For this purpose, we profiled the whole-blood transcriptome of two native breeds and one commercial breed using high-throughput RNA sequencing. More than 15,000 genes were identified, of which 2, 89, and 162 genes were significantly upregulated exclusively in Northern Finncattle, Yakutian cattle and Holstein cattle, respectively. The functional classification of these significantly differentially expressed genes identified several biological processes and pathways related to signalling mechanisms, cell differentiation, and host-pathogen interactions that, in general, point towards immunity and disease resistance mechanisms. The gene expression pattern observed in Northern Finncattle was more similar to that of Yakutian cattle, despite sharing similar living conditions as the Holstein cattle included in our study. In conclusion, our study identified unique biological processes in these breeds that help them to adapt and survive in sub-arctic environments.

## Introduction

As a result of natural and human-led selection, domestic animals have been able to adapt, survive, be productive and reproduce in challenging environments (Mirkena et al., 2010). Animals in the northern hemisphere (Lapland, northern Russia and Siberia) are used for food production and other socio-cultural needs and, thus, have been the basis of human life in the North. These animals may have different biological capacities to adapt to extremes in temperature, daylight and feed availability. However, there has been growing interest towards improving local breeds by introducing genetic material from superior breeds, with a preference towards commercial breeds. Intensive breeding and the replacement of native breeds with commercial breeds may appear to be advantageous at first, but these practices will have long-lasting consequences. In addition to irreversibly losing the unique genetic resources of native breeds, we will also be losing the cultural and historical heritage associated with those local breeds. The signatures of adaptation, as well as the history of formation, that are encoded in the native breeds may also fade away (Gaouar et al., 2015). Therefore, the local breeds must be genetically characterized and preserved.

Here, we have studied two native cattle breeds (Northern Finncattle and Yakutian cattle) and one international breed (Holstein cattle). All of these breeds are used for milk and meat production. Previous studies using genetic markers have indicated the genetic distinctiveness among these breeds (Li et al., 2007; Li and Kantanen, 2010). Among the three breeds, Holstein cattle have high economic importance, due to high productivity, and have the shortest adaptation history. Holstein cattle are the most popular dairy breed globally, with an intensive selection programme and high milk production; although they originated in a temperate climate, they have adapted to different parts of the world and can survive in varying climatic conditions. Northern Finncattle and Yakutian cattle have economic, social and cultural values (Kantanen et al., 2015). The Northern Finncattle breed is native to Northern Finland and Finnish Lapland. The breed nearly became extinct during the1970s but is currently maintained (current census is 850 cows) through active *in vivo* and *in vitro* conservation activity. The Yakutian cattle are characterized by being purebred aboriginal native cattle. Adult Yakutian cows (current population is approx. 1,000 animals) typically weigh 350-400 kg, and their height is 111 cm, on average. The animals are well adapted to harsh Siberian conditions, where the temperature falls below −50°C in long winters (Kantanen et al., 2009).

High-throughput RNA sequencing (RNA-Seq) has been proven to be an efficient method for studying the regulation of gene expression (Wang et al., 2009). In recent years, RNA-Seq has been applied to domestic animals, with a major focus on productivity traits (Bai et al., 2016; Li et al., 2016; Pokharel et al., 2018; Silva-Vignato et al., 2017). A number of transcriptome studies have been conducted in different Holstein breeds (Bai et al., 2016; Li et al., 2016; Sandri et al., 2015), but to date, there are no reports on gene expression studies in either Yakutian cattle or Northern Finncattle. In this study, we have applied RNA-Seq technology and profiled the whole blood transcriptome of the three aforementioned breeds to characterize their genetic differences.

## Materials and methods

### Sample collection

Animal handling procedures and sample collection were conducted in accordance with legal regulations that were approved by the Russian authorization board (FS/UVN-03/163733/07.04.2016) and the Animal Experiment Board in Finland (ESAVI/7034/04.10.07.2015). A 2.5-ml sample of blood from each of three Yakutian cattle, three Northern Finncattle and three Holstein cattle cows was collected into a PAXgene® Blood RNA IVD tube (Ref. PreAnalytiX®, Hombrechtikon, Switzerland) during the winter period of 2016 and 2017 and stored at −18°C. The Finnish samples were taken in a slaughterhouse at exsanguination, and the Yakutian samples were collected by jugular venipuncture in the northern Yakutian village Sakkyryr (YC6) and central Yakutia Magaras (YC10 and YC11). All of the animals included in this study were 4- to 8-year-old females, except for one 14-year-old Holstein cow (HC3).

### RNA extraction

The RNA was extracted with the PAXgene® Blood RNA kit (Ref. 762174, Ref. PreAnalytiX®, Hombrechtikon, Switzerland), according to the kit manual, with minor adjustments to the protocol: the samples were thawed at room temperature overnight, the initial centrifugation time at 5,000 × g was increased to 15 minutes, the initial pellet was resuspended into double the amount of BR1 buffer and divided into two separate reactions, the proteinase K incubation time was extended to 1 hour, and the columns were incubated at room temperature in elution buffer for 10 minutes prior to elution. The concentration and quality of the RNA was measured with a spectrophotometer (NanoDrop ND-1000, Thermo Fisher Scientific, Delaware, USA), and the integrity of the RNA was measured with an Agilent Bioanalyzer 2100 (Agilent, Waldbronn, Germany) using the Agilent 6000 RNA Nano kit (Ref. 5067-1511, Agilent, Waldbronn, Germany). All samples selected for sequencing had an RNA Integrity Number (RIN) value of at least 7.

### Library preparation and sequencing

Library preparation and sequencing tasks were outsourced to the Finnish Functional Genomics Center (FFGC) in Turku, Finland. The library preparation was performed according to Illumina’s Truseq® mRNA sample preparation guide protocol. The high quality of the libraries was confirmed with an Advanced Analytical Fragment Analyzer, and the concentrations of the libraries were quantified using Qubit® Fluorometric Quantitation, Life Technologies. The average RNA-Seq library fragments were in the range of 250-350 bp. Only the good libraries with RNA Quality Number (RQN) value greater than 7 were sequenced with a paired-end strategy to generate 75 bp reads.

### Computational methods

Raw sequence reads were pre-processed using FastQC (Simon Andrews, n.d.) to determine the quality of the data and to obtain an overview of the sequencing data. The outputs from FastQC were summarized using Multiqc (Ewels et al., 2016). Processed reads were mapped against the latest versions of the cattle reference genome UMD3.1 and transcriptome (Ensembl release 93) using STAR v2.6 (Dobin et al., 2013). First, the genome indices were prepared, and mapping was performed with default parameters. Moreover, a gene-level counts file for each sample was generated as part of the star-alignment pipeline. Read counts were then processed using DESeq2 (Love et al., 2014) for gene expression analysis, and all genes that had less than 5 transcript counts were discarded. To identify significantly differentially expressed genes between breed groups, we set an adjusted p-value of 0.05. For the list of significantly differentially expressed genes, we retrieved additional information, such as gene description and chromosomal location, using the Biomart Bioconductor package (Durinck et al., 2009; Smedley et al., 2009). Finally, the significantly differentially expressed genes were assessed for their functional roles. Gene ontology (GO) terms and the Kyoto Encyclopaedia of Genes and Genomes (KEGG) pathway analyses were performed using the ClueGO (Bindea et al., 2009) plugin in Cytoscape (v3.6.1) (Shannon et al., 2003). In addition to the default parameters in ClueGO, we chose to observe GO terms in the range of level 3 and 5. Similarly, for the GO term or KEGG pathway to be displayed, at least three genes and a minimum of 4% of the total genes should be present in our list. In addition, the terms or pathways shared by 50% of the genes from our list were grouped together using Kappa statistics (Kappa score threshold of 0.4).

## Results and discussion

From nine samples, we obtained 32.2 Gb of RNA-Seq data. More than 87% of the reads from each sample mapped to the cattle reference genome, with Holstein samples having more than 91% mapping rates. The slightly higher mapping rate for Holstein samples could be because that breed is closer to the Hereford breed, which is the source of the reference assembly (Zimin et al., 2009). With sequencing costs becoming cheaper, having breed-specific reference genomes would be useful in the future.

A total of 15,960 genes (Table S1) were expressed in our data, which covers 64.8% of known (24,616) cattle genes reported in the annotation file. At the individual level, we observed that both the highest (FC8) and the lowest (FC7) number of genes were expressed in Northern Finncattle samples (Table 1). We believe that a comparatively lower number of reads in the sample FC7 led to a lower number of expressed genes. We did not observe any globin genes among the top most expressed genes. Genes such as MHC class I heavy chain (BOLA), uncoupling protein 2 (UCP2), eukaryotic translation elongation factor 3 (EEF2) and vimentin (VIM) were among the top most expressed genes. These results confirmed that RNA extraction using PAXgene® worked well.

**Table 1:** Sample summary.

| Sample ID | Breed | ENA Accession | No. of reads (million) | Uniquely mapped reads % | No. of genes expressed |
| --- | --- | --- | --- | --- | --- |
| FC7 | Northern Finncattle | ERS2639473 | 49.4 | 87.2 | 12,106 |
| FC8 | Northern Finncattle | ERS2639474 | 107.6 | 87.1 | 15,038 |
| FC9 | Northern Finncattle | ERS2639475 | 89.4 | 89.7 | 14,811 |
| HC1 | Holstein | ERS2639476 | 104.4 | 91.7 | 14,768 |
| **HC2** | Holstein | ERS2639477 | 62.8 | 91.0 | 14,140 |
| **HC3** | Holstein | ERS2639478 | 62.4 | 92.4 | 13,831 |
| **YC6** | Yakutian | ERS2639479 | 69.2 | 87.3 | 14,047 |
| **YC10** | Yakutian | ERS2639480 | 69.8 | 88.2 | 13,736 |
| **YC11** | Yakutian | ERS2639481 | 65.8 | 88.5 | 14,371 |

We assessed the co-expression status of the top 500 most expressed genes (i.e., the genes with highest base mean values) in each of the three breed groups. As seen in the Venn diagram (Fig. 1), 386 of the top expressed genes were present in all three breeds. Holstein samples had the highest number (n = 65) of unique genes, followed by Yakutian cattle (n = 56) and Northern Finncattle (n = 17). Moreover, the results showed that Northern Finncattle are more closely related to Yakutian cattle (53 uniquely shared genes) than to Holstein cattle (44 uniquely shared genes). In contrast, Yakutian cattle and Holstein cattle shared the least number of unique genes.

**Fig. 1:**
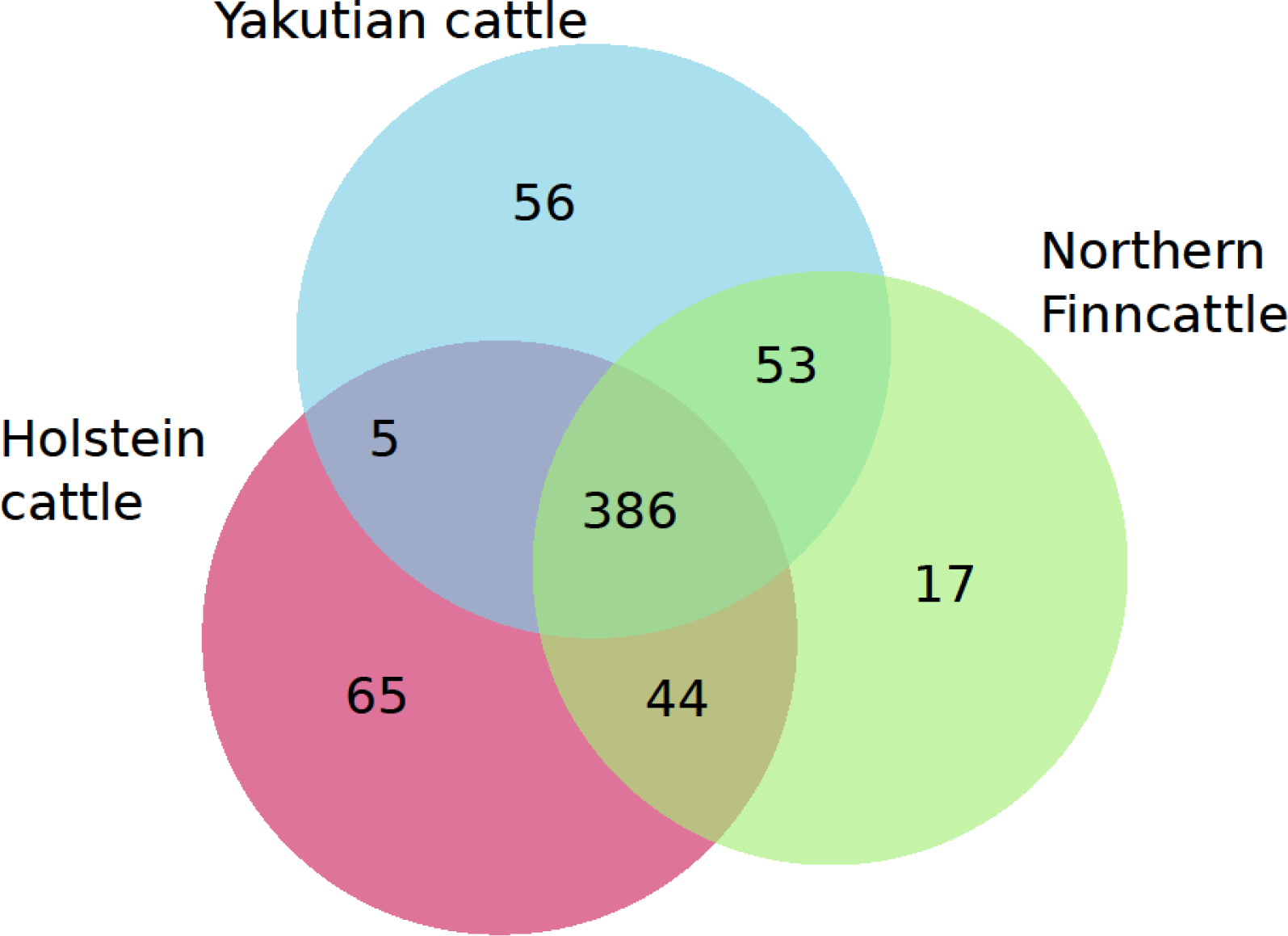
Co-expression of the top 500 genes among the three breed groups.

We then looked more closely into the genes that were differentially expressed between any two of the three breed groups and assessed what types of biological processes and/or biological pathways those genes were associated with.

### Differential gene expression between Yakutian cattle and Northern Finncattle

A total of 189 transcripts were significantly differentially expressed between Yakutian cattle and Northern Finncattle (Table S2), of which 108 transcripts were upregulated in Yakutian cattle.

Transcripts of known genes, such as *C-C motif chemokine ligand 5* (*CCL5*), *perforin 1* (*PRF1*), *SLAM family member 7* (*SLAMF7*), *CD300 molecule-like family member b* (*CD300LB*), *sphingosine-1-phosphate receptor 5* (*S1PR5*) and *ATP binding cassette subfamily A member 13* (*ABCA13*), were upregulated in Yakutian cattle. Genes upregulated in Northern Finncattle included *transglutaminase 3* (*TGM3*), *5’-aminolevulinate synthase 2* (*ALAS2*), *ATP synthase inhibitory factor subunit 1* (*ATP5IF1*) and *FK506 binding protein 5* (*FKBP5*). Moreover, a number of genes with unknown functions were upregulated in Yakutian cattle (e.g., ENSBTAG00000047816, ENSBTAG00000038141, ENSBTAG00000010300, and ENSBTAG00000046323) and in Northern Finncattle (e.g., ENSBTAG00000046981, ENSBTAG00000048145, ENSBTAG00000003754, and ENSBTAG00000002450). Having a better annotation would certainly help in characterizing the functions of these genes. The GO terms associated with the upregulated genes in Yakutian cattle included “positive regulation of natural killer cell chemotaxis”, “regulation of phagocytosis, negative regulation of viral process”, “negative regulation of cytokine production” and “granzyme-mediated apoptotic signalling pathway” (Fig. 2a, Table S3). The KEGG pathways associated with upregulated genes included “antigen processing and presentation”, “natural killer cell mediated cytotoxicity” and “graft-versus-host disease” (Table S4). Similarly, the GO terms associated with upregulated genes in Northern Finncattle included “regulation of lipid transport”, “erythrocyte differentiation”, “regulation of cytokine-mediated signalling pathway” and “regulation of protein maturation” (Fig. 2b, Table S5). “Malaria” (associated genes *HBA*, *HBB* and *THBS1*) was the only KEGG pathway related to genes upregulated in Northern Finncattle.

**Fig. 2:**
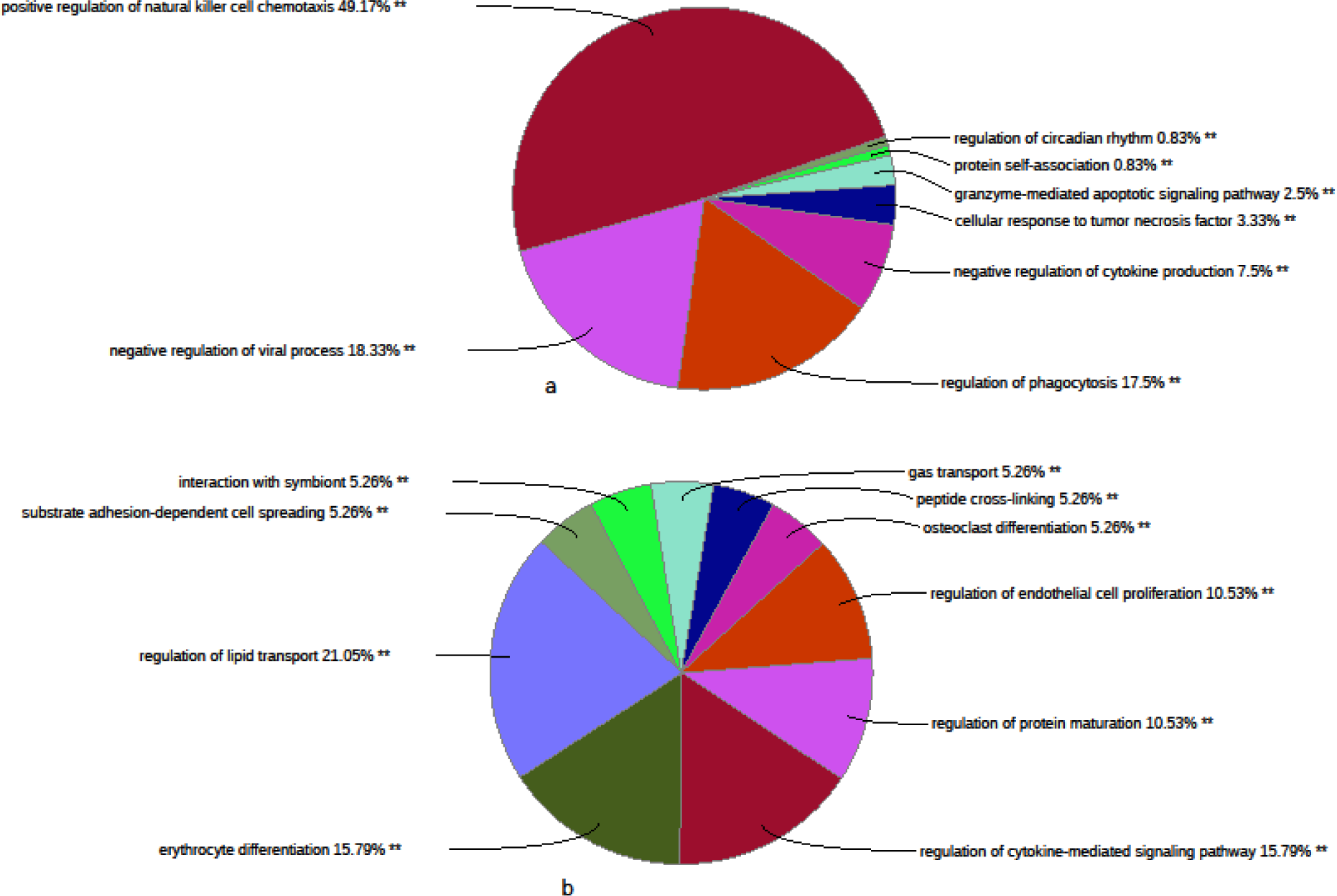
A functional annotation of the significantly differentially expressed genes in Yakutian cattle and Northern Finncattle. The GO terms associated with upregulated genes in Yakutian cattle (a) and Northern Finncattle (b). Similar GO terms were grouped together, and thus, the percentages at the ends of each representative GO term indicate the total percentages of linked GO terms. The significant GO terms at the p<0.001 statistical level are indicated by double (**) asterisks.

### Differential gene expression between Yakutian cattle and Holstein cattle

The highest number of significantly differentially expressed genes was identified between Yakutian cattle and Holstein cattle (Table S6). Given the phenotypic difference between the breeds, it was not surprising to observe a large number of differentially expressed genes. Yakutian cattle have phenotypic characteristics, such as a small and compact body size and a thick hair coat, which helps them survive in a cold environment. In contrast, Holstein cattle have a bigger body size and a thin hair coat and are thus not ready for a colder climate. In addition, Holstein cattle do not face a shortage of fodder and are maintained inside the barn, whereas Yakutian cattle may suffer from food shortages and are partly outside during the winter. The significant differences between Yakutian cattle and Holstein cattle are in line with earlier reports (Decker et al., 2016; Yurchenko et al., 2018; Zinovieva et al., 2016). Out of 1,418 genes that were significantly differentially expressed between the two breeds, 594 genes were upregulated in Yakutian cattle. Because of the high number of differentially expressed genes, we only considered those genes with a log2-fold change greater than or less than 1.5 for GO and KEGG pathway analyses. Genes that were upregulated in Yakutian cattle were associated with GO terms such as “chemokine-mediated signalling pathway”, “regulation of natural killer cell-mediated immunity”, “kinase activator activity” and “regulation of T cell activation” (Fig. 3a, Table S7). In contrast, the genes that were upregulated in Holstein cattle were associated with GO terms such as “inflammatory response”, “regulation of neutrophil chemotaxis”, “regulation of response to external stimulus” and “secondary active transmembrane transporter activity” (Fig. 3c, Table S8). The KEGG pathways associated with upregulated genes in Yakutian cattle included “natural killer cell-mediated cytotoxicity”, “African trypanosomiasis”, “endocrine and other factor-regulated calcium reabsorption”, “cytosolic DNA-sensing pathway”, “cell adhesion molecules (CAMs)” and “cytokine-cytokine receptor interaction” (Fig. 3b, Table S9). Similarly, pathways such as “non-small cell lung cancer”, “legionellosis”, “notch signalling pathway”, “cytokine-cytokine receptor interaction”, “adipocytokine signalling pathway”, “osteoclast differentiation”, “protein digestion and absorption” and “ABC transporters” were related to the genes that were upregulated in Holstein cattle (Fig. 3d, Table S10).

**Fig. 3:**
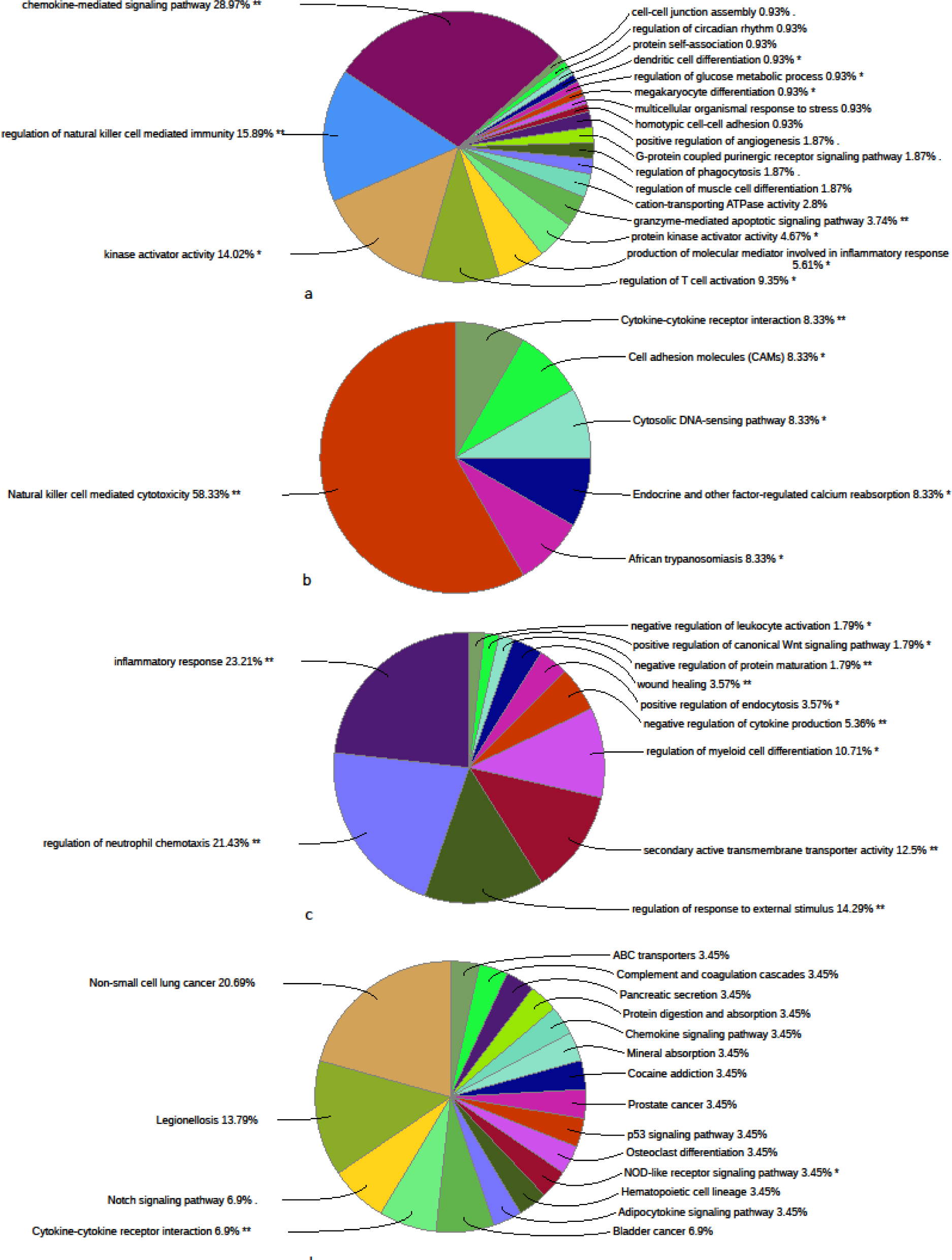
A functional annotation of the genes that were significantly differentially expressed between Yakutian cattle and Holstein cattle. (a) The GO terms and (b) KEGG pathways that were associated with upregulated genes in Yakutian cattle. (c) The GO terms and (d) KEGG pathways that were associated with upregulated genes in Holstein cattle. Similar GO terms and KEGG pathways were grouped together, and thus, the percentages at the ends of each term indicate the total percentages of linked GO terms and KEGG pathways. The significant GO terms and KEGG pathways at the p<0.05 and p<0.001 statistical levels are indicated by single (*) and double (**) asterisks, respectively.

### Differential expression between Northern Finncattle and Holstein cattle

Between Finncattle and Holstein cattle, we observed 250 significantly differentially expressed transcripts (Table S11), with majority the of genes (n = 180) being upregulated in Holstein cattle. Only one GO term, “eicosanoid metabolic process” (associated genes *ALOX15*, *ALOX5*, *HPGD*), and no KEGG pathways were identified for genes that were upregulated in Northern Finncattle. In contrast, GO terms, such as “response to molecule of bacterial origin”, “cellular response to lipopolysaccharide”, “negative regulation of haemopoiesis”, and “response to mechanical stimulus” (Fig. 4, Table S12), and KEGG pathways, such as “osteoclast differentiation”, “complement and coagulation cascades”, “TNF signalling pathway” and “Toll-like receptor pathway”, were associated with upregulated genes in Holstein cattle (Table S13).

**Fig. 4:**
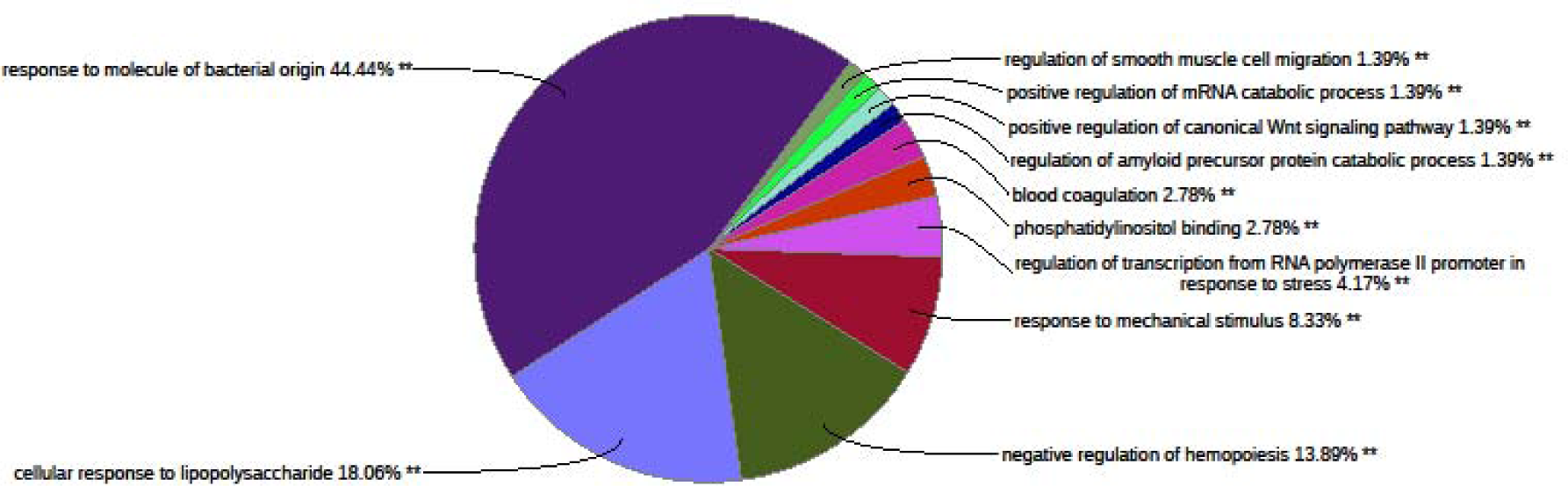
A functional annotation of the genes that were upregulated in Holstein cattle compared to Northern Finncattle. Similar GO terms were grouped together, and thus, the percentages at the ends of each term indicate the total percentages of linked GO terms. The significant GO terms at the p<0.001 statistical level are indicated by double (**) asterisks.

### Uniquely differentially expressed genes

We noticed that there was overlap among many genes in more than one comparison (Fig. S1). Therefore, we compiled a list of genes that were uniquely differentially expressed in each breed. The number of uniquely differentially expressed genes (Table 2) in each breed followed a similar pattern to that observed by the top 500 most expressed genes (Fig. 1).

**Table 2:** Numbers of uniquely differentially expressed genes in different breeds.

| **Breed** | **No. of uniquely differentially expressed genes** | **Upregulated genes** | **Downregulated genes** |
| --- | --- | --- | --- |
| **Yakutian cattle** | 139 | 89 | 52 |
| **Northern Finncattle** | 9 | 2 | 7 |
| **Holstein cattle** | 200 | 162 | 48 |

Yakutian cattle had 139 uniquely differentially expressed genes, of which 89 were reported to be upregulated. We noticed a number of cases where more than one gene from the same family was upregulated. Such cases could provide higher confidence related to gene functions. The genes or receptors with more than one member are as follows: chemokines (*CCL4*, *CCL5*), carbohydrate sulfotransferases (*CHST1*, *CHST12*), chemokine receptors (*CX3CR1*, *CXCR6*), growth arrests (*GAS6*, *GAS7*), granzymes (*GZMB* (two paralogs), *GZMM* and *GZMH*), insulin-like growth factor binding proteins (*IGFBP4*, *IGFBP7*) and natural cytotoxicity triggering receptors (*NCR1*, *NCR3*). Among the list were *BHLHE40* and *PRKCG*, which are linked to circadian rhythm. The importance of the circadian clock is particularly applicable to Yakutian cattle. They must be metabolically prepared for food scarcity during long winters (Ebling and Barrett, 2008).

Four granzyme transcripts and perforin were upregulated in Yakutian cattle. Granzymes are serine proteases that are used by cytotoxic lymphocytes to destroy malignant and virus-infected cells. Granzymes are transported into the cytoplasm of the target cell by *perforin 1* (*PRF1*), after which they cleave specific proteins and trigger apoptosis (Johnson et al., 2003; MacDonald et al., 1999; Russell and Ley, 2002). It has been suggested that the evolution of granzymes is related to species-specific immune challenges (humans have 5 granzyme genes and mice have 10), which they maintain by gene duplications and alterations in substrate specificity (Kaiserman et al., 2006). Three out of the six granzymes known in cattle, (Yang et al., 2018) with two nearby paralogs of *granzyme B* (Fig. S2) and *PRF1*, are all upregulated in Yakutian cattle. These findings suggest that Yakutian cattle have a very solid granzyme-mediated immune system.

The number of upregulated genes was sufficient to identify significant GO terms. The GO terms associated with the upregulated genes included “negative regulation of cytokine production”, “regulation of lymphocyte-mediated immunity”, “positive regulation of leukocyte chemotaxis” and “cellular response to interferon-gamma” (Fig. 5A, Table S14). Similarly, pathways such as “natural killer cell-mediated cytotoxicity”, “inflammatory bowel disease (IBD)”, “African trypanosomiasis” and “cytosolic DNA-sensing pathway” were related to uniquely upregulated genes in Yakutian cattle (Fig. 5b, Table S15). Only one GO term, “regulation of lipid transport” (associated genes *ABCA1*, *ABCG1*, *IRS2* and *THSB1*), and no KEGG pathways were found for the downregulated genes.

**Fig 5:**
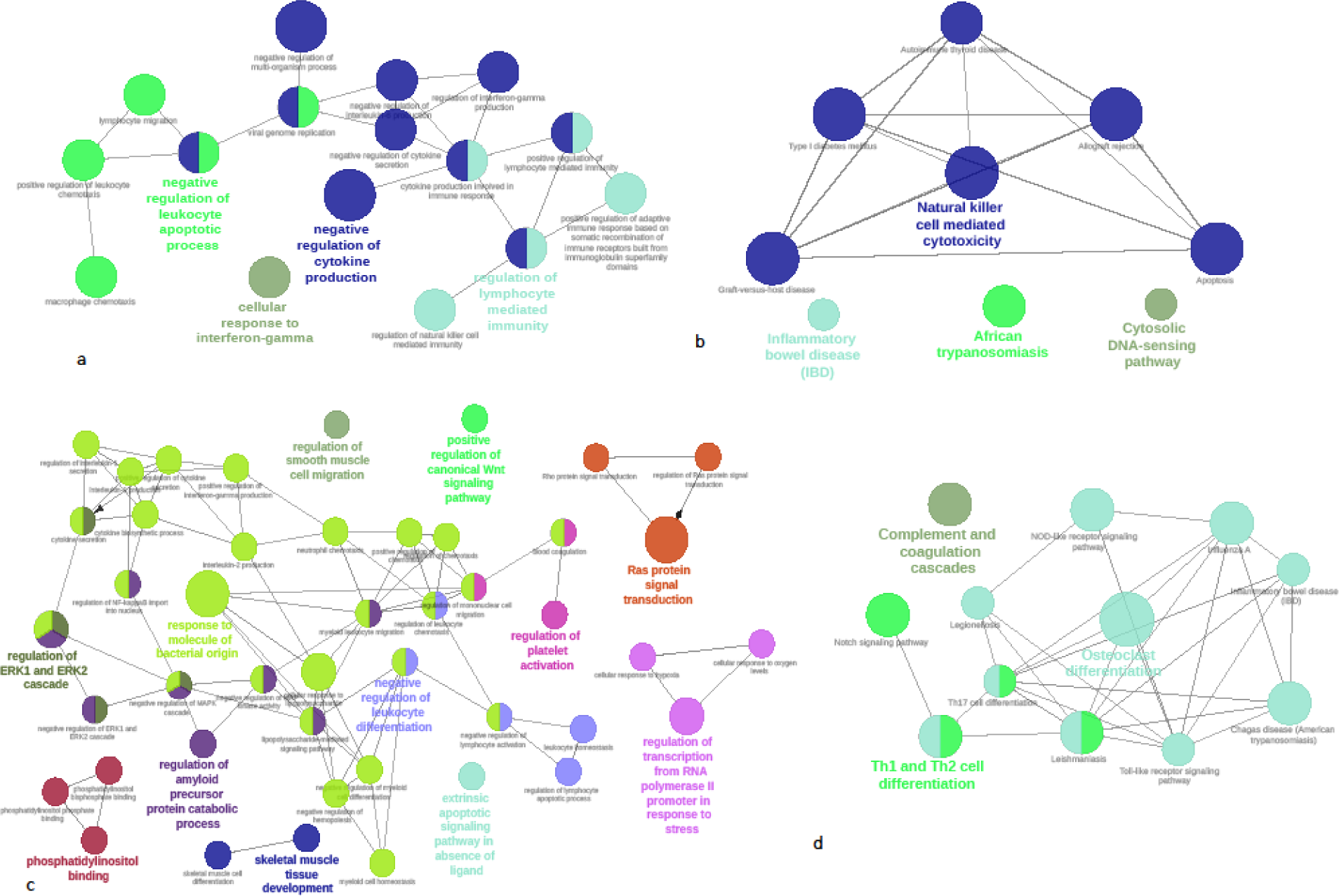
A functional annotation of the significantly differentially expressed genes between Yakutian and Holstein cattle. Nodes represent GO terms, with node size corresponding to the significance of term enrichment, and functionally related groups partially overlap. A network of (a) GO terms and (b) KEGG pathways that are associated with the genes upregulated in Yakutian cattle. A network of (c) GO terms and (d) KEGG pathways that are associated with upregulated genes in Holstein cattle. The representative GO terms and KEGG pathways are highlighted in bold.

The majority (162/200) of the uniquely differentially expressed genes in Holstein cattle were found to be upregulated. The list also includes a number of cases where more than one member of the gene family was present. Genes encoding the solute carrier protein family (*SLC16A12*, *SLC16A6*, *SLC25A37*, *SLC28A3*, *SLC2A9*, *SLC45A4*, *SLC6A6*, *SLC23A1*), ring finger proteins (*RNF149*, *RNF24*), notch proteins (*NOTCH1*, *NOTCH2*, *NOTCH3*), lysine demethylases (*KDM4B*, *KDM6B*), interferon receptors (*IFNAR1*, *IFNAR2*, *IFNGR1*), CD proteins (*CD101*, *CD164*, *CD300a*, *CD55*), and others were upregulated in Holstein cattle samples. The GO terms associated with the upregulated genes included “cellular response to lipopolysaccharide”, “response to molecule of bacterial origin”, “negative regulation of leukocyte differentiation”, “regulation of transcription from RNA polymerase II promoter in response to stress and phosphatidylinositol binding” (Fig. 5C, Table S16). Similarly, KEGG pathways such as “osteoclast differentiation”, “Th1 and Th2 cell differentiation” and “complement and coagulation cascades” were related to the upregulated genes (Fig. 5d, Table S17). We did not find any GO terms or KEGG pathways associated with the downregulated genes.

Among other functions, membrane-bound solute carriers (SLCs) are key to maintaining physiological processes, including nutrient uptake, waste removal and ion transport (Hediger et al., 2004). The upregulated expression of eight SLC proteins in Holstein cattle samples suggest the highly important function of these genes. A recent study showed that, during sloughing, amphibians increase the rate of ion uptake to maintain internal homeostasis (Wu et al., 2017). We speculate that, with a comparatively bigger body size and a thin hair coat, Holstein cattle may have developed a similar system for maintaining internal homeostasis in response to lower temperatures. The upregulation of three notch proteins (*NOTCH1*, *NOTCH2* and *NOTCH3*) and the biological processes associated with hypoxia (“cellular response to oxygen levels”, “cellular response to hypoxia”) is in line with earlier findings, where the “notch signalling pathway” was activated in response to hypoxia (Hiyama et al., 2011). This finding also suggests that Holstein cattle may not have adapted well to winter conditions, as hypoxia is linked to low temperatures (Rocha and Branco, 1998). “Notch signalling pathway” is also known to have important role in milk lactose metabolism, mammary gland development and lactation which are all associated with high milk production traits of Holstein cattle (Politi et al., 2004; Yalcin-Ozuysal et al., 2010). Moreover, *NOTCH1* and *NOTCH2* are part of the processes “Ras protein signal transduction” and “regulation of ERK1 and ERK2 cascade”.

*Keratin 72* (*KRT72*, ENSBTAG00000007904) and *transmembrane protein 8 A* (*TMEM8A*, ENSBTAG00000016588) were the only genes that were exclusively upregulated in Northern Finncattle. *KRT72* is involved in hair formation, and the function of *TMEM8A* is unknown. Out of seven downregulated genes, four IDs (ENSBTAG00000038233, ENSBTAG00000039691, ENSBTAG00000046611, ENSBTAG00000000930) do not have gene descriptions available. The remaining three genes are *acetylcholinesterase* (*ACHE*), *chemokine (C-C motif) ligand 3* (*CCL3*) and *Bos taurus regulator of G-protein signalling 2, 24 kDa* (*RGS2*). The RGS family of proteins is involved in fine-tuning the signalling activities of G-protein-coupled receptors (GPCRs) (Berman et al., 1996; Blumer, 2004). An increased level of *RGS2* expression is linked to the regulation of numerous biological activities, such as immune responses, bone formation, cardiovascular function and anxiety (Zhang and Mende, 2014). *CCL3*, *CCL4* and *CCL5* are members of the CC chemokine family of proteins, which exhibit proinflammatory activities. Moreover, *CCL3* inhibits the proliferation of haematopoietic stem/progenitor cells (HSPCs) (Cook, 1996). *ACHE* has numerous functions, and its role in muscle development, neuritogenesis, cell adhesion, the activation of dopamine neurons, and amyloid fibre assembly has been reviewed earlier (Soreq and Seidman, 2001).

### Limitations of the study

Finally, we would like to highlight some of the limitations of this study that we are aware of. First, having more samples could give higher confidence to the results. Comparisons involving samples from temperate climates would be equally interesting. Due to the quality of the gene annotations, many genes could not be functionally interpreted. Moreover, due to the large number of differentially expressed genes, we did not look in detail at all of the genes, and therefore, many interesting genes may not have been highlighted in this study. While we tried to interpret our results based on model species, primarily human studies, some of the interpretation may not be accurate. Particularly, most of the studies are based on model species with more controlled settings, whereas the animals in our study, especially Yakutian animals, live in harsh environmental conditions.

## Conclusions

In conclusion, using blood as a starting material, we were able to capture more than 60% of cattle genes to assess the differential gene expression in three breeds. The gene expression profiles in two important indigenous cattle breeds that are adapted to the northern and even sub-arctic climate have been reported for the first time. Gene expression comparisons between the three breed groups revealed that Northern Finncattle are more closely related to Yakutian cattle than to Holstein cattle, despite sharing similar living conditions with the Holstein cows that were analysed in this study. Moreover, we identified breed-specific differences in maintaining immunity and disease resistance mechanisms among the breeds included in our study. Some of our results, based on Holstein cattle, explain how alternative biological processes can help imported animals cope with challenging environmental conditions. Studies such as this one could identify genetic markers that may assist in animal breeding and the sustainable utilization and conservation practices of animal genetic resources in changing northern Eurasian environments.

## Data availability

The raw sequence reads (FASTQ files) have been deposited to the European Nucleotide Archive (ENA) under the accession number PRJEB28074 (please refer to Table 2 for sample specific accessions). The script used during the differential expression analysis is available as supplementary code (Code S1).

## Acknowledgements

We thank our colleague Tuula Marjatta Hamama from the Natural Resources Institute Finland (Luke) for laboratory assistance and Innokenty Ammosov for valuable assistance in sampling in Yakutia. This study belongs to the Arctic-Ark project, funded by the Academy of Finland (decision no. 286040). The authors wish to acknowledge the CSC – IT Center for Science, Finland, for computational resources. This study was supported by the Finnish Functional Genomics Centre, University of Turku and Åbo Akademi and Biocenter Finland. Special thanks to the owners of the experimental animals for letting us collect the blood samples.

## Author contributions

J.K. conceived the analysis. J.K., M.H., T.R., J.P., H.L., R.P. and S.Z. participated in sample collection. H.H. extracted mRNA from blood samples. K.P. and M.W.B. performed RNA-Seq data analyses. K.P. wrote the manuscript. All authors read, revised and approved the final manuscript. The authors declare that they have no competing interests.

## Supplementary files

Fig. S1: Venn diagram based on the list of significantly differentially expressed genes following all three possible comparisons

Fig. S2: Figure showing the closely located Granzyme B in the genome Table S1: Table listing the expression levels of each sample

Table S2: List of significantly differentially expressed genes between Yakutian cattle and Northern Finncattle

Table S3: The GO terms associated with upregulated genes in Yakutian cattle when compared with Northern Finncattle

Table S4: The KEGG pathways associated with upregulated genes in Yakutian cattle when compared with Northern Finncattle

Table S5: The GO terms associated with upregulated genes in Northern Finncattle when compared with Yakutian cattle

Table S6: Summary of significantly differentially expressed genes between Yakutian cattle and Holstein cattle

Table S7: The GO terms associated with upregulated genes in Yakutian cattle when compared with Holstein cattle

Table S8: The GO terms associated with upregulated genes in Holstein cattle when compared with Yakutian cattle

Table S9: The KEGG pathways associated with upregulated genes in Yakutian cattle when compared with Holstein cattle

Table S10: The KEGG pathways associated with upregulated genes in Holstein cattle when compared with Yakutian cattle

Table S11: Summary of significantly differentially expressed genes between Northern Finncattle and Holstein cattle

Table S12: The GO terms associated with upregulated genes in Holstein cattle when compared with Northern Finncattle

Table S13: The KEGG pathways associated with upregulated genes in Holstein cattle when compared with Northern Finncattle

Table S14: The GO terms associated with the genes that were exclusively upregulated in Yakutian cattle

Table S15: The KEGG pathways associated with the genes that were exclusively upregulated in Yakutian cattle

Table S16: The GO terms associated with the genes that were exclusively upregulated in Holstein cattle

Table S17: The KEGG pathways associated with genes that were exclusively upregulated in Holstein cattle

Code S1: R script that was used to identify and annotate differentially expressed genes in this study

